# Degradation of benzene by the heavy-metal resistant bacterium *Cupriavidus metallidurans* CH34 reveals its catabolic potential for aromatic compounds

**DOI:** 10.1101/164517

**Authors:** Felipe A. Millacura, Franco Cárdenas, Valentina Mendez, Michael Seeger, Luis A. Rojas

## Abstract

Benzene, toluene, ethylbenzene and the three xylene isomers are monoaromatic contaminants widely distributed on polluted sites. Some microorganisms have developed mechanisms to degrade these compounds, but their aerobic and anaerobic degradation is inhibited in presence of heavy metals, such as mercury or lead. In this report, the degradation of benzene and other aromatic compounds catalyzed by the metal resistant bacterium *Cupriavidus metallidurans* CH34 was characterized. A metabolic reconstruction of aromatic catabolic pathways was performed based on bioinformatics analyses. Functionality of the predicted pathways was confirmed by growing strain CH34 on benzene, toluene, *o-*xylene, *p-*cymene, 3-hydroxybenzoate, 4-hydroxybenzoate, 3-hydroxyphenylacetate, 4-hydroxyphenylacetate, homogentisate, catechol, naphthalene, and 2-aminophenol as sole carbon and energy sources. Benzene catabolic pathway was further characterized. Results showed that firstly benzene is transformed into phenol and, thereafter, into catechol. Benzene is degraded under aerobic conditions via a combined pathway catalyzed by three Bacterial Multicomponent Monooxygenases: a toluene-2-monoxygenase (TomA012345), a toluene-4-monooxygenase (TmoABCDEF) and a phenol-2-hydroxylase (PhyZABCDE). A catechol-2,3-dioxygenase (TomB) expressed at early exponential phase cleaves the catechol ring in *meta*-position; an *ortho*-cleavage of catechol is accomplished by a catechol-1,2-dioxygenase (CatA) at late exponential phase instead. This study additionally shows that *C. metallidurans* CH34 is capable of degrading benzene in presence of heavy metals, such as Hg(II) or Pb(II). This capability of degrading aromatic compounds in presence of heavy metals is rather unusual among environmental bacteria; therefore, *C. metallidurans* CH34 seems to be a promising candidate for developing novel bioremediation process for multi-contaminated environments.

**HIGHLIGHTS:**

1. The strain *Cupriavidus metallidurans* CH34 is capable to degrade benzene aerobically
2. Benzene oxydation is mediated by bacterial multicomponent monoxygenases
3. Strain CH34 is able to grow using a broad range of aromatic compounds as sole carbon and energy source
4. Benzene degradation occurs even in presence of heavy metals such as mercury and lead

## Introduction

Monoaromatic molecules such as benzene, toluene, ethylbenzene and the three-xylene isomers (*ortho*, *meta* and *para*) are commonly known as BTEX (Parales et al., 2008; Choi et al., 2013). They are part of the volatile fraction of petroleum hydrocarbons and are often found in industrial polluted sites as remnant of chemical products, fuels, solvents or lubricants (Fuchs et al., 2011; Fuentes et al., 2014). Their toxicity is well known due to their mutagenic and carcinogenic effects exerted via bioaccumulation in animal and human tissues (Browning, 1961; Dean, 1978; Fuentes et al., 2014). The most hazardous and toxic BTEX is benzene which causes cancer and leukimia in humans (Dean, 1978; World Health Organization, 1993; van der Park, 2014), currently considered as the fourth priority substance in the environmental quality standards upheld within the European Union (EU Parliament, 2008). Benzene not only contaminates soils but also ground water and atmosphere (Browning, 1961; Lovley, 1995). The low level of benzene permitted on potable water in the United States demonstrates that it is considered a high risk for human health. Indeed, US maximum levels for BTEX in potable water are 0.05, 1.00, 0.70 and 10 ppm for benzene, toluene, ethylbenzene and the xylene isomers, respectively (USEPA, 2006).

Although these aromatic compounds are toxic, some bacteria have developed mechanisms to survive in contaminated environments using these compounds as substrate for their growth (Fuchs et al., 2011; Fuentes et al., 2014). For instance, Benzene is oxidized into phenol by the bacterial multicomponent monooxygenases (BMM) present in *C. pinatubonensis* JMP134 or into *cis*-benzenediol catalyzed by the benzene dioxygenase from *P. putida* F1 (Reardon et al., 2000). Both pathways converge in the central intermediate catechol that is thereafter cleaved and degraded through the tricarboxylic acid cycle (Zamanian and Mason, 1987; Bertoni et al., 1998). Therefore, the use of microorganisms arises as a promising strategy for the clean-up of aromatic compounds, such as petroleum hydrocarbons, pesticides and chlorophenols. Successful examples of *in situ* soil bioremediation performed by bacteria have been described (Chen et al., 2015). However, either aerobic and anaerobic degradation of BTEX is inhibited in sites co-contaminated with heavy metals, such as mercury and lead (Kovalick, 1991; Muniz et al., 2004; Davydova, 2005; Kavamura and Esposito, 2010; Dórea et al., 2007).

Heavy metals and BTEX compounds are widespread together in the environment due to diverse anthropogenic factors, e.g., urban and mining activities. Mercury has been extensively used in gold amalgam extraction, whereas lead has been used for many decades in its tetraethyl form as a fuel additive (Veiga and Meech, 1991; Nascimento and Chartone-Souze, 2003; Sandrin and Maier, 2003; Seyferth, 2003; Kovarik, 2005; Kristensen et al., 2014). As BTEX contamination is predominantly originated by oil- and petroleum spills, a third of sites contaminated with organic compounds are also contaminated with inorganic compounds (Kovalick, 1991). In fact, approximately 40% of the hazardous waste sites in the US are contaminated simultaneously with organic and inorganic contaminants (Sandrin and Maier, 2003). Additionally, European Union (2008) stipulated benzene, lead, and mercury, as respectively the fourth, twentieth, and twenty-first priority substances in terms of European environmental quality standards (EU Parliament, 2008). As environmental problems remain a major challenge, development of novel bioremediation approaches to assess bioremediation of co-contaminated sites is urgently required.

Bacterial resistance to heavy metals is well-documented (Silver, 1996; Mergeay et al., 2003; Smalla et al., 2006; Rojas et al., 2011; Altimira et al., 2012). Particularly, the strain *Cupriavidus metallidurans* CH34 is a heavy metal-resistant model bacterium that harbors two large plasmids, pMOL28 and pMOL30, which carry genetic determinants for heavy metal resistance (Mergeay et al., 1985; Mergeay et al., 2003; Monchy et al., 2007; Janssen et al., 2010). Furthermore, diverse catabolic clusters have been detected on the genome of *C. metallidurans* strains and *Cupriavidus* sp. (Mergeay et al., 1985; Janssen et al., 2010; Pérez-Pantoja et al., 2012; Rosier et al., 2012; Mergeay and van Houdt, 2014; Basu et al., 2016), although its actual degrading potential has not yet been assessed. This research characterizes the degradation pathways of benzene and other aromatic compounds presents on the metal resistant bacterium *C. metallidurans* CH34. In addition, the growing of *C. metallidurans* CH34 on benzene in presence of mercury and lead was examined, in order to forecast future applications on bioremediation of co-contaminated sites.

## Materials and Methods

### Chemicals

Benzene, toluene, phenol and *o*-xylene (>99.7% purity) were obtained from Merck (Darmstadt, Germany), 3-hydroxybenzoate, 4-hydroxybenzoate, 3-hydroxyphenylacetate, 4-hydroxyphenylacetate, 2,5-dihydroxyphenylacetate (homogentisate), catechol, 2-aminophenol, naphthalene and *p*-cymene (>98% purity) were obtained from Sigma Aldrich (St. Louis, MO, USA). HgCl_2_ and Pb(NO_3_)_2_ were obtained from Merck (Darmstadt, Germany) and used to prepare Hg(II) and Pb(II) stock solutions. Sodium succinate dibasic hexahydrate was obtained from Sigma (Steinheim, Germany; >99.0% purity).

### Bacterial strains and culture conditions

*C. metallidurans* CH34, *Pseudomonas putida* mt-2 and *Pseudomonas putida* G7 were cultivated in low-phosphate Tris-buffered mineral salts (LPTMS) medium at 30°C. The LPTMS medium contained (per 1 L): 6.06 g Tris Base USP (US Biological, Swampscott MA, USA); 4.68 g NaCl (Merck, Darmstadt, Germany); 1.07 g NH_4_Cl (Merck, Darmstadt, Germany); 1.49 g KCl (Merck, Darmstadt, Germany); 0.43 g Na_2_SO_4_ (Merck, Darmstadt, Germany); 0.2 g MgCl_2_•6H_2_O (J.T. Baker, Phillipsburg, NJ, USA); 0.03 g CaCl_2_•H_2_O (Merck, Darmstadt, Germany); 0.005 g Fe(III)(NH4) citrate (Merck, Darmstadt, Germany), and 1 mL of trace element solution SL7 of Biebl and Pfennig (Mergeay et al., 1985; Rojas et al., 2011). Additionally, *Burkholderia xenovorans* LB400 was cultivated at 30°C in mineral M9 medium with an elemental trace solution (Méndez et al., 2011). Succinate (10 mM), benzene (5 mM), or other aromatic compounds (1 mM) were used as sole carbon and energy source, provided directly in liquid phase if not otherwise stipulated. Growth assays were performed in triplicate measuring turbidity at 600 nm. To determine the Hg(II) and Pb(II) Minimal Inhibitory Concentrations (MICs) during growth of CH34 on benzene, the bacteria were grown on liquid LPTMS minimal medium using benzene (5 mM) as only carbon and energy source. Bacteria were challenged to increasing concentrations of Hg(II) and Pb(II) from stock solutions of HgCl_2_ and PbCl_2_ analytical grade (Sigma Aldrich Saint Louis, MO, USA).

### Bioinformatic analysis

Genome of *C. metallidurans* CH34 has been sequenced and mostly annotated (Janssen et al., 2010). Sequences for chromosome (NC_007973.1), chromid (NC_007974.2), pMOL28 (NC_007972.2) and pMOL30 (NC_007971.2) were obtained from the GenBank database. The metabolic reconstruction was based on standard protocols (Thiele and Palsson, 2010; Nogales, 2014). An initial draft was generated using SEED 2.0 (Overbeek et al., 2005). Predictions were refined and curated manually by applying NCBI/BLAST searches (http://blast.ncbi.nlm.nih.gov/Blast.cgi), the metabolic database MetaCyc (http://metacyc.org/), and the database of genes and genomes of Kyoto, KEGG (http://www.genome.jp/kegg/). Prediction of promoter regions was performed using BacPP software (de Avila e Silva, 2011) augmented by protein association analysis using STRING v9.1 software (Franceschini et al., 2013). Orthologous gene sequence analysis was performed at aminoacid level using Clustal-W software (http://www.ebi.ac.uk/Tools/msa/clustalw2/) under default parameters. In addition, the organization of gene clusters involved in BTEX degradation for CH34 with other BTEX degradative bacteria was compared by SEED viewer 2.0 tool. Metabolic pathway images were generated using the Ultra ChemBioDraw 13.0 software from Perkin Elmer.

### RNA Isolation

Total RNA was isolated from *C. metallidurans* CH34 using the RNeasy mini kit (Qiagen, Hilden, Germany) according to the manufacturer’s recommendations. TURBO DNAfree set (LifeTechnologies, Carlsbad, USA) was used to degrade any residual DNA. A final qPCR test with *gyrB* primers designed by Primer 3.0 (Table S1) was performed in order to confirm a total degradation of DNA. The RNA concentration was quantified using a Qubit fluorometer (Invitrogen) and a Nanodrop spectrophotometer (Thermo Scientific). RNA integrity was tested by agarose (1%) gel electrophoresis.

### Real-Time RT-PCR

Reverse transcription was carried out using 200 ng of RNA and was achieved with a High Capacity cDNA Reverse Transcription Kit (Applied Biosystems, California, USA). The Minimum Information for publication of Quantitative real-time PCR Experiments (MIQE) guideline was used as standard protocol (Taylor et al., 2010). Real-time PCR was performed using 20 ng of cDNA on a StepOne Real-Time PCR System (Applied Biosystems, California, USA), using Maxima SYBR Green/ROX qPCR Master Mix (Thermo Scientific, California USA) and 0.3 μM of each primer. cDNA was initially denatured at 95°C for 5 min. A 40-cycle amplification and quantification protocol (95°C for 15 s, 55°C for 15 s and 60°C for 15 s) with a single fluorescence measurement per cycle followed by a melting-curve program (95°C for 15 s, 25°C for 1 s, 50°C for 15 s and 95°C for 1 s) were used according to the manufacturer’s recommendations. PCR melting curves confirmed the amplification of a single product for each primer pair. Primers yielded products between 200-250 bp. The *gyrB* (Rmet_0003) gene was amplified as a reference gene, yielding an amplicon of 233 bp. A standard curve in triplicate was made with serial dilutions (10 fold) for each amplicon in a linear range (10 ng – 0.1pg) of genomic DNA. qPCR efficiencies were calculated from the slopes of the log-linear portion of calibration curves, using the equation E=10 ^(1/slope)^. Reference *gyrB* gene was stably expressed according to the algorithms of BestKeeper (Pfaffl et al., 2004). Relative gene expression ratios were determined as outlined by Pfaffl in 2001 (Pfaffl, 2001), thereby normalizing gene expression levels of CH34 cells grown on benzene versus CH34 cells grown on succinate.

### Intermediates detection

Aliquots were taken at different times during the growth of *C. metallidurans* CH34 on benzene (5mM). Cells were lysed by sonication, centrifuged (19,000 × *g* for 5 min) and cell-free supernatants analyzed using a Jasco high performance liquid chromatograph (HPLC) model LC-2000 equipped with a diode array detector (DAD) Jasco model MD-2015 plus a RP 18e/Chromolith column of 100-4.6 mm (Merck, Darmstadt, Germany). The solvents used for sample elution were 0.1% formic acid in water (A) and 100% acetonitrile (B). The flow rate was 1.0 mL/min and the elution profile was 70% A:30% B for 4 min, then changed linearly to 0% A:100% B over a 1 min period and kept at this ratio for 3 min and finally changed linearly to 30% A:70% B over a 1 min period and kept at this ratio for 2 min. Benzene and phenol were quantified using calibration curves with authentic standards. Experiments were performed in triplicate. The formation of 2-hydroxymuconic semialdehyde (HMS) was determined during growth using a Perkin Elmer Lambda UV/VIS spectrophotometer by measuring the absorbance at 375 nm. HMS concentrations were calculated using the molar extinction coefficient of catechol as previously described (Nozaki et al., 1970).

## Results

### Genomic analysis of CH34 genes involved in benzene degradation

Genes encoding benzene peripheral and central catabolic pathways are located in two different chromosomal clusters. A first locus with a size of 24,547 bp is comprised from Rmet_1305 to Rmet_1331 (Table S2), and a second locus with an extension of 7,587 bp, including from Rmet_1781 to Rmet_1788 (Table S3). Both clusters encode in total three Bacterial Multicomponent Monooxygenases (BMMs), two catechol dioxygenases (C23O and C12O), two transcriptional regulators (XylR/NtrC-type), a membrane transport protein (TbuX/FadL-type) and ten enzymes part of the central metabolism of diverse aromatics (Figure 1).

**Figure 1.**
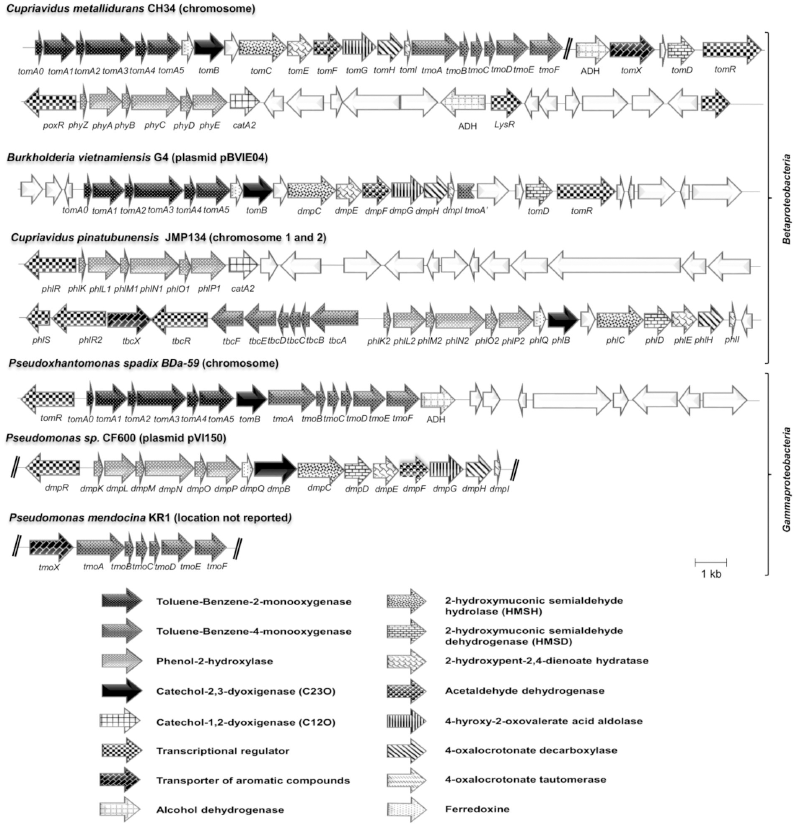
Organization of gene clusters involved in the catabolism of benzene in *C. metallidurans* CH34 and other *Proteobacteria*. The orientations of ORFs are represented by open arrows. Sizes of genes and intergenic regions are to scale.

Interestingly, the gene clusters encoding monooxygenases from strain CH34 share a high synteny with aromatic degradative clusters from different *Proteobacteria* (Figure 1, Table S2). For instance, the organization of genes (*tmoABCDEF*) encoding the toluene-4-monooxygenase (T4MO) in *C. metallidurans* CH34, as well as the aa sequences of their products, are highly similar to the layout of the corresponding gene clusters and encoded product sequences from *C. pinatubonensis* JMP134 (TBC), *Pseudoxanthomonas spadix* BD-a59 (TMO), *Pseudomonas mendocina* KR1 (TMO) and *P. stutzeri* OX1 (TBU). The cluster (*phyZABCDE*) encoding the phenol-2-hydroxylase (PHY) from strain CH34 shows a high similarity in gene organization and amino acid (aa) sequence with the gene clusters from *C. pinatubonensis* JMP134 (PHL) and *Pseudomonas* sp. CF600 (DMP). The toluene-2-monooxygenase (T2MO) from strain CH34, encoded on *tomA012345*, presents a high similarity with the corresponding loci from *B. vietnamiensis* G4 (T2MO) and *Pseudoxanthomonas spadix* BD-a59 (T2MO). Finally, the predicted *tomBCEFGHI* gene cluster that encode enzymes of the central catabolic pathway in *C. metallidurans* CH34 is highly similar in organization and sequence to the *tomB*, *dmpCEFGHI*, *tomD* and *tomR* genes from *B. vietnamiensis* G4 and *dmpBCDEFGHI* from *Pseudomonas sp*. CF600 (pVI150) (Figure 1, Table S2).

Both entire likely to be regulated by the XylR/NtrC-type transcriptional regulators TomR (Rmet_1305) and PoxR (Rmet_1788), highly related with their orthologs present on *B. vietnamiensis* G4 (97% aa) and *C. pinatubonensis* JMP134 (90% aa), respectively (Table S3). Additionally, the *tomX* (Rmet_1326) gene encodes a TbuX/FadL-type membrane transport protein that possesses high similarity with the transporter TbuX from *B. multivorans* DDS15A-1, which is part of the Toluene_X superfamily of monoaromatic outer membrane transport proteins (Hearn et al., 2009). We further identified the presence transport proteins for other aromatic compounds. An ABC transporter permease for benzoate that is similar to BenK (Nishikawa et al., 2008), encoded formerly on genes located from Rmet_1226 to Rmet_1230, and a protocatechuate and 4-hydroxybenzoate transporter that is part of the superfamily of mayor facility transporters PcaK, encoded formerly on gene Rmet_4011 (Harwood et al., 1994; Janssen et al., 2010).

### **Growth of *C. metallidurans* CH34 on aromatic compounds**

This study reveals that *C. metallidurans* CH34 is capable of growing in liquid LPTMS minimal medium using benzene, toluene, *o-*xylene, *p-*cymene, 3-hydroxybenzoate, 4-hydroxybenzoate, 3-hydroxyphenylacetate, 4-hydroxyphenylacetate, homogentisate, catechol, naphthalene or 2-aminophenol as sole carbon and energy source (Table 1). Reaching and tolerating concentrations similar to model bacteria such as *P. putida* mt-2 (toluene degrader), *P. putida* G7 (naphthalene degrader) and *B. xenovorans* LB400 (biphenyl degrader), used in this study as reference strains for toluene, naphthalene, and biphenyl degradation, respectively. Although strain CH34 is capable of growing on diverse aromatics, it was not able to use other aromatic compounds as metabolic energy source, such as bisphenol A, phenanthrene, anthracene, vanillate, nitrobenzene, *m*-toluic acid, 4-isopropylbenzoic acid and 1,2,4-benzenetriol.

**Table 1:** Growth of *C. metallidurans* CH34 on aromatic compounds as sole carbon and energy sources

| Carbon source | *C. metallidurans* CH34 | *B. xenovorans* LB400 | *P. putida* mt-2 | *P. putida* G7 |
| --- | --- | --- | --- | --- |
| benzene<sup>a</sup> | ++ | NT | + | - |
| toluene<sup>a</sup> | +++ | NT | +++ | - |
| *o*-xylene<sup>a</sup> | + | NT | - | - |
| *p*-cymene<sup>a</sup> | +++ | NT | + | + |
| 3-hydroxybenzoate | ++ | + | +++ | +++ |
| 4-hydroxybenzoate | +++ | +++ | +++ | +++ |
| 3-hydroxyphenylacetate | ++ | ++ | +++ | +++ |
| 4-hydroxyphenylacetate | +++ | +++ | ++ | ++ |
| naphtalene | ++ | + | - | +++ |
| 2-aminophenol | ++ | ++ | NT | NT |
| homogentisate | +++ | +++ | +++ | +++ |
| catechol | +++ | +++ | +++ | +++ |
| succinate or glucose<sup>b</sup> | +++ | +++ | +++ | +++ |

Growth level observed (+) slight, (++) moderate, (+++) high; (-) no growth; (NT) not tested.
<sup>a</sup>Compounds provided in gaseous phase.
<sup>b</sup>For comparison purposes, bacteria were grown on succinate (strains CH34, G7 and mt-2) or glucose (strain LB400), and mentioned as positive just if values obtained overpassed strain growth without carbon source.

Interestingly, strain CH34 is able to tolerate benzene concentrations up to the saturation point in water (20 mM) and is able to grow on presence of xylene isomers (*o- m- p*-) mixes (data not shown). Even though the growth on *p*-cymene, xylene (*o- m- p*-) isomers, 3-hydroxybenzoate, naphthalene, and 2-aminophenol was observed, orthologous genes for *xyl, cym/cum*, *nah*, *amn* were not found among the genome of CH34.

### Metabolic intermediates during benzene degradation

In order to analyze the functionality and metabolic intermediates formation of the benzene pathway, growth assays on benzene were performed (Figure 2A). A yellow colorization was observed after 20 h of growth on benzene, which disappeared after 48 h. The color change of the growing culture suggests an active *meta-*cleavage pathway, likely due to the formation of 2-hydroxymuconic semialdehyde (Nozaki et al., 1970). In order to identify the metabolic intermediates generated during the growth, culture supernatants of CH34 cells grown on benzene (5 mM) were analyzed by HPLC (see Material and Methods). The results showed that benzene concentration decreased over time, while the appearance of phenol was observed after 15-26 h of growth (Figure 2B). Phenol is known to be an intermediate on benzene oxidation catalyzed by T2MO, T3MO and T4MO (Nozaki et al., 1970). Phenol is then transformed into catechol and further converted into intermediates from the catabolic central pathways through *meta* or *ortho* ring cleavage. Catechol was not detected by HPLC, maybe due to it fast degradation. Formation of 2-hydroxymuconic semialdehyde (HMS) was observed after 22h (early exponential phase), which suggests catechol degradation through *meta*-cleavage (Figure 2B). The formation of this intermediate occurs at an early exponential phase. Suggesting that *meta*-cleavage of the catechol ring is catalyzed by a catechol-2,3-dioxygenase (TomB) during early states of growth.

**Figure 2.**
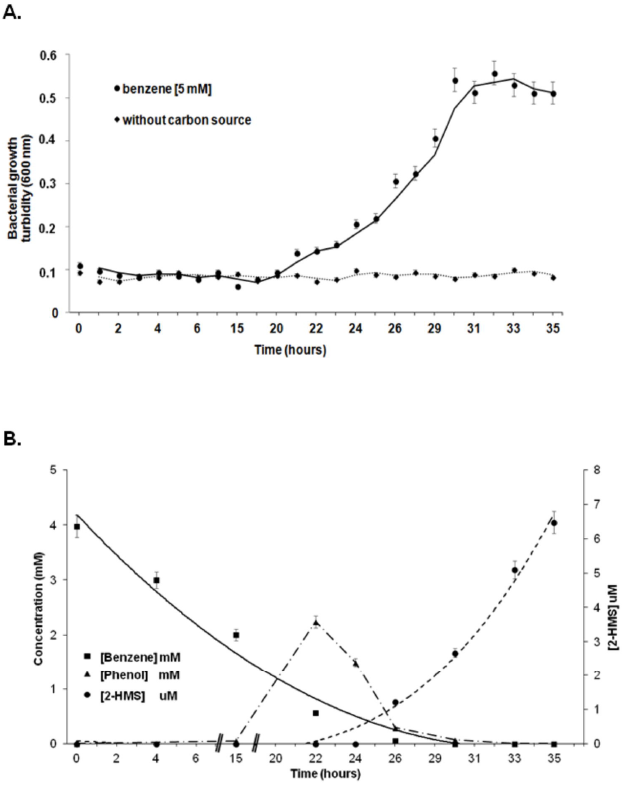
Formation of the metabolic intermediates phenol and semialdehyde-2-hydroxymuconic during the growth of *C. metallidurans* CH34 on benzene. A. CH34 cells were grown in LPTMS minimal medium using benzene (5 mM) as sole carbon and energy sources. Control assay without carbon source are also depicted. **B**. The metabolic intermediates were analyzed by HPLC. Benzene degradation (squares), phenol formation (triangle) and the generation of semialdehyde-2-hydroxymuconic (2-HMS; circle) after *meta*-cleavage of the catechol ring are indicated. Control assays without bacteria showed no degradation (data not shown). Each point is an average ± SDs of results from at least three independent assays.

### Transcriptional analysis during benzene degradation

Formation of 2-hydroxymuconic semialdehyde (HMS) was observed in cells grown on benzene during the exponential phase (Figure 2B); therefore, CH34 cells were grown on benzene (5 mM) and collected at early (turbidity of 0.2~0.3) and late (turbidity of 0.5~0.6) exponential phase. The expression of genes encoding the monooxygenases TOM (*tomA3*), TMO (*tmoA*), PHY (*phyC*), the dioxygenases C12O (*catA1* and *catA2*) and C23O (*tomB*), and the enzymes HMSD (*tomC*) and HMSH (*tomD*), were quantified. In addition, the transcription of the sigma*-*38 factor gene (*rpoS*) and the LysR-type (*catM* and *benM*) as well as the XylR/NtrC-type (*tomR* and *poxR*) transcriptional regulators were studied. Real time RT-PCR analysis showed a simultaneous expression of genes encoding monooxygenases in both early and late exponential phases (Figure 3). The *tomB* (C23O), *catA1* (C12O) and *catA2* (C12O) genes display a differential expression. *tomb* gene was induced in the early and late exponential phases, whereas, *catA1* and *catA2* genes showed a low expression in the early stage of growth, and a further induction in the late exponential phase. In addition, the *tomC* and *tomD* genes were also expressed in early and late exponential phases showing that both pathways of HMS are active. The *tomR* gene encoding a XylR/NtrC-type transcriptional regulator was expressed in early and late exponential phases. In contrast, the sigma*-*38 factor gene (*rpoS*) and the *benM, catM,* and *poxR* genes encoding transcriptional regulators are expressed only in the late exponential phase (Figure 3).

**Figure 3.**
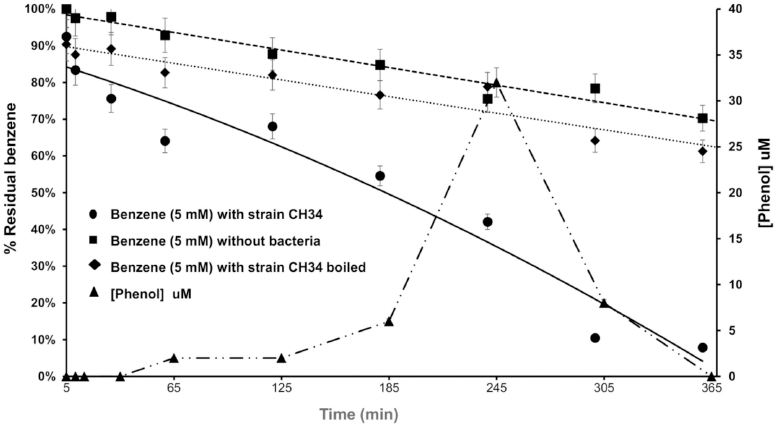
Transcriptional analysis of genes from the benzene catabolic pathway. RT-qPCR assays were performed using mRNA from CH34 cells grown on LPTMS minimal medium supplemented with benzene (5 mM) until early exponential phase (turbidity at 600 nm of 0.2~0.3; dark grey column) and late exponential phase (turbidity at 600 nm of 0.5~0.6; grey column). The genes encode for toluene-benzene-2-monooxygenase (*tomA3*), toluene-4-monooxygenase (*tmoA*), phenol-2-monooxygenase (*phyC*), catechol-2,3-dioxygenase (*tomB*), catechol-1,2-dioxygenase (*catA1* and *catA2*), hydroxymuconic semialdehyde dehydrogenase (*tomC*), 2-hydroxymuconic semialdehyde hydrolase (*tomD*), sigma factor 38 (*rpoS*), LysR-type transcriptional regulators (*benM* and *catM*), XylR/NtrC-type transcriptional regulators (*tomR* and *poxR*). The *gyrB* gene was used as a reference gene. The primer pairs used are listed in Table S1. The fold-change in gene expression was calculated relative to CH34 cells grown in succinate. *p* value= 0.1%.

### **Growth of *C. metallidurans* CH34 on benzene in presence of heavy metals**

Strain CH34 was capable of growing on benzene in presence of mercury concentrations up to 0.005 mM (~1 ppm). Likewise, the MIC to Pb(II) for strain CH34 was only slightly affected when benzene was used as a sole carbon source and its growth was unaffected at Pb(II) concentration of up to 0.2 mM (~82 ppm). The MIC was recorded as the lowest concentration (mM) of Hg(II) and Pb(II) at which no growth was observed (Table 2).

**Table 2:** Minimal inhibitory concentration of Hg(II) and Pb(II) for CH34 cells grown on succinate or benzene.

| Metal [mM] | Strain CH34 | Strain CH34 | Ratio<br>(fold)<sup>a</sup> |
| --- | --- | --- | --- |
|  | Succinate | Benzene |  |
| Hg<sup>+2</sup> | 0.025 | 0.008 | -3.0 |
| Pb<sup>+2</sup> | 0.600 | 0.400 | -1.5 |

<sup>a</sup>Ratio (fold) expressed as heavy metal resistance from CH34 cells grown on succinate compared to cells grown on benzene 5 mM.

## Discussion

In this study, we report metabolic insights of benzene degradation by *C. metallidurans* CH34 in order to understand the capabilities of strain CH34 to degrade aromatic compounds, even in the presence of heavy metals. Metabolic reconstruction of aerobic benzene degradation was performed based on genomic analysis, gene expresion, intermediates detection, transcriptional analysis and growth studies.

Two aerobic benzene catabolic pathways have been described (Bertoni et al., 1998; Reardon et al., 2000; Tao et al., 2004). Benzene can be oxidized by a BMM into phenol or by a benzene dioxygenase into *cis*-benzenediol (Zamanian and Mason, 1987; Bertoni et al., 1998). *C. metallidurans* CH34 possesses chromosomal gene clusters encoding three BMMs (Notomista et al., 2003; Janssen et al., 2010). The function of these BMMs associated to the degradation of benzene and other aromatic compounds was shown in this study (Figure 3). The findings obtained from the genomic studies reveal that BMM toluene-2-monooxygenase (T2MO) subunits (encoded by *tomA012345* genes) possess a high similarity in amino acid sequence and gene organization to the T2MO subunits from *B. vietnamiensis* G4 and *P. spadix* BD-a59 (Figure 1 and Table S2), suggesting a regiospecific hydroxylation of toluene into *o-*cresol and, subsequently, an oxidation into 3-methylcatechol (Shields et al., 1989; Hur et al., 1997; O’Sullivan et al., 2007). In addition, T2MO catalyzes the oxidation of dichloroethylenes, chloroform, 1,4-dioxane, aliphatic ethers, and diethyl sulphide (Hur et al., 1997; Ryoo et al., 2000; Mahendra and Alvarez-Cohen, 2006), and enables the formation of epoxides from a variety of alkene substrates (McClay et al., 2000). On the other hand, the toluene-4-monooxygenase protein (T4MO) encoded by the *tmoABCDEF* gene cluster is similar to the enzymes from *P. spadix* BD-a59, *P. mendocina* KR1 and *P. pnomenusa* 3kgm (Figure 1 and Table S2). The T4MO from strain KR1 oxidizes toluene into 4-methylcathecol and catalyzes the formation of epoxides from a variety of alkene substrates (McClay et al., 2000), as well as catalyzes the oxidation of phenols and methylphenols into catechol (Shields et al., 1989). Additionally, the T4MO from *P. stutzeri* OX1 has the capability to oxidise *o*-xylene, *m*-xylene, *p*-xylene, toluene, benzene, ethylbenzene, styrene, naphthalene and tetrachloroethylene (Hur et al., 1997). Overall, T2MO and T4MO monooxygenases are capable of catalyzing three successive hydroxylations on benzene to form phenol, catechol and 1,2,3-trihydroxybenzene, respectively (Tao et al., 2004).

Located downstream of the *tomA012345* gene cluster, the gene cluster *tomBCEFGHI* encodes enzymes for the *meta*-cleavage and the subsequent reactions from the central pathway of benzene degradation. These enzymes are similar in amino acid sequence to the gene products of corresponding gene clusters present in *B. vietnamiensis* G4 and *Pseudomonas* sp. CF600 (Figure 1 and Table S2). The findings obtained from the genomic studies also suggest that bacteria carrying the *dmp-*encoded central pathway from *Pseudomonas* sp. CF600 share the BMM DmpKLMNOP. However, the strain CH34 possesses a gene organization that includes a different BMM upstream (T2MO) and downstream (T4MO) of this central catabolic pathway (Figure 1). Downstream of the genes encoding the benzene central catabolic pathway from CH34, a cluster of genes that encode a toluene-4-monooxygenase (*tmoABCDEF*) is located. The products of these genes are similar to the corresponding subunits from strains *C. pinatubonensis* JMP134 (TBC), *P. mendocina* KR1 and *P. spadix* BD-a59 (TMO) (Figure 1). All these bacteria have different BMMs, such as toluene monooxygenases permitting not only degrading benzene and phenol but also other BTEX compounds (Tao et al., 2004), as is the case in strain CH34. Based on results obtained from genomic analysis of genes that are involved in transport and degradation of benzene and aromatic compounds in strain CH34, we propose novel metabolic pathways for the aerobic degradation of aromatic compounds in *C. metallidurans* CH34 (Figure 4). Strain CH34 seems to have different routes for catalyzing successive hydroxylations to convert benzene into phenol and catechol as proposed in Figure 4. In strain CH34, these gene clusters are likely regulated by the *tomR* gene product (formerly Rmet_1305), which is a XylR/NtrC-type transcriptional regulator (Table S2).

**Figure 4.**
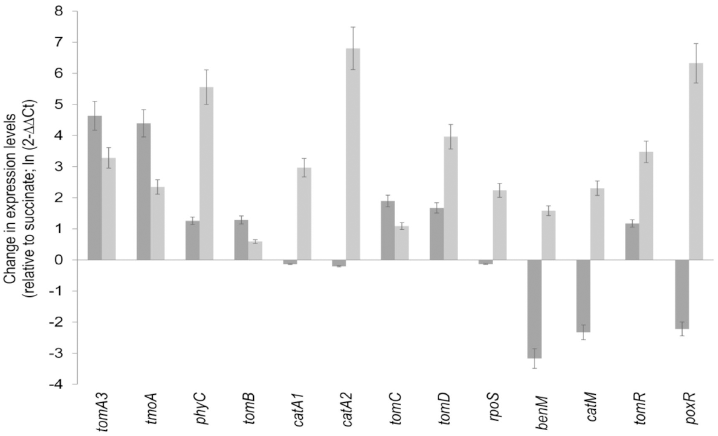
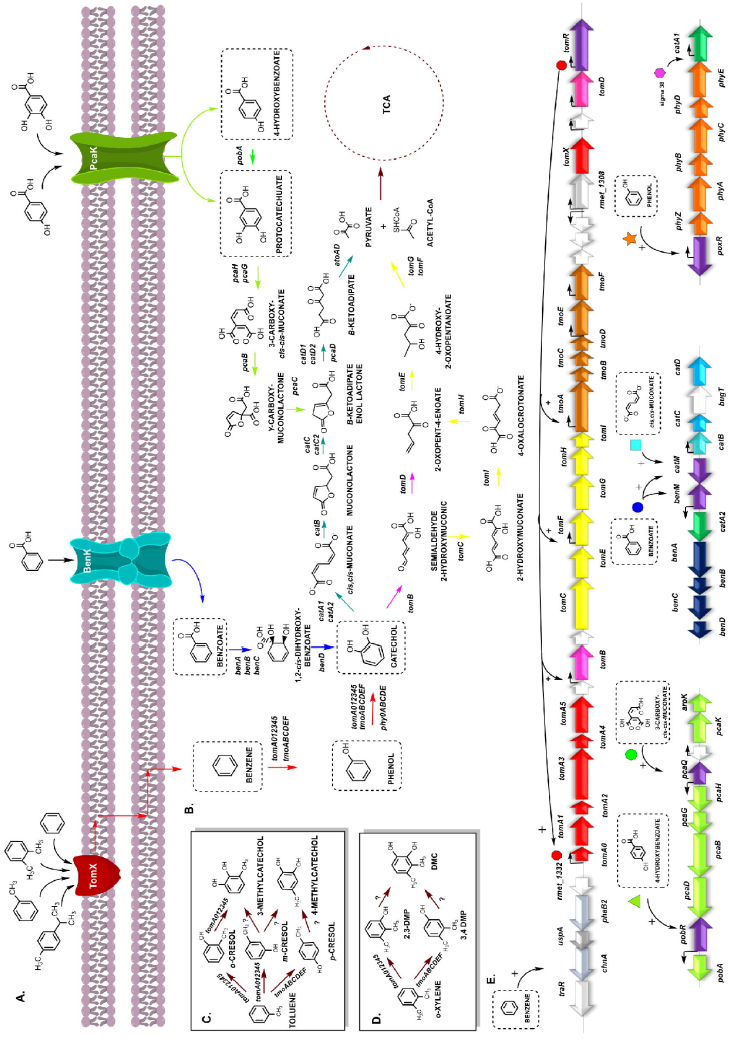
Model of aerobic aromatic compounds degradation in *C. metallidurans* CH34. Gene(s), substrate(s) and product(s) of each enzyme are indicated. **A**. Transporters: monoaromatic hydrocarbons FadL/TbuX-type transporter (TomX, red); benzoate ABC-type transporter (BenK, blue); protocatechuate and 4-hydroxybenzoate belonging to the Major Facilitator Superfamily (PcaK, green). **B**. Peripheral and central catabolic pathways catalyzed by toluene-2-monooxygenase (*tomA012345*, red arrow), phenol-2-hydroxylase (*phyZABCDE*, orange arrow) and toluene-4-monooxygenase (*tmoABCDEF*, brown arrow), benzoate-1,2-dioxygenase (*benAB*, blue arrow), 1,2-*cis*-dihydroxybenzoate dehydrogenase (*benD*, blue arrow), 4-hydroxybenzoate monooxygenase (*pobA,* green arrow), C23O (*tomB*, pink arrow), C12O (*catA1* and *catA2*, green arrow), HMSH (*tomD,* pink arrow); enzymes from the central catabolic pathways for monoaromatic compounds (*tomCEFGHI*, yellow arrow), catechol (*catBCD*, light blue arrow), protocatechuate (*pcaBCDGH*, green arrows) and entrance to the tricarboxylic acid cycle (TCA, dark red dotted line). **C**. Organization and proposed gene regulation: LysR-type transcriptional regulators (purple arrow), CatM (blue square), BenM (blue circle) and PcaQ (green hexagon); AraC type transcriptional regulator (PobR, green triangle); XylR/NtrC-type transcriptional regulators dependent of *sigma* 54 factor (early exponential phase) TomR (red hexagon) and PoxR (orange star). Benzene is detected by TomR and triggers activation of peripheral degradation pathway (*tmoABCDEF* and *tomA012345*). Phenol is detected by TomR and triggers a conformational change and activation of the central degradation pathway (*tomBCEFGHID*); PoxR recognizes presence of phenol in the system and activates *phyZABCDE* genes. The *sigma* 38 factor (purple hexagon) present at late exponential growth phase regulates the transcription of the C12O *catA2* located downstream from the *phy* genes. Presence of *cis*,*cis*-muconate is recognized by CatM and BenM generating a synergistic activation of the *ortho*-catechol degradation pathway. The entry of 4-hydroxybenzoate to the cell is recognized by PobR triggering a transcriptional activation of the *pobA* gene. Formation of 3-carboxy-*cis*,*cis*-muconate generates expression of the central protocatechuate degradation pathway. The promoter regions are denoted with small black arrows bent in the directions of transcription and were identified by BacPP. Protein-protein interactions were analyzed using STRING v9.1 software. The sizes of genes and intergenic regions are to scale.

The CH34 gene cluster *phyZABCDE* encodes the third BMM, a phenol-2-hydroxylase (P2MO). This cluster is also present in *C. pinatubonensis* JMP134 and *W. numazuensis* TE26 (Janssen et al., 2010). Downstream of the *phyZABCDE* gene cluster from CH34 is located a *catA2* gene that encodes a catechol-1,2-dioxygenase (C12O) (Figure 1 and Table S3). A second gene encoding for a C12O is located on the chromid (Rmet_4881) and belongs to the benzoate degradation pathway (Perez-Pantoja et al., 2012). A previous study has reported that the C12O enzyme from strain CH34 is unique in its capacity to cleave diverse catechols in *ortho* position, *e.g*. tetrachlorocatechol, 4-fluorocatechol, 4-methylcatechol, and 3-methylcatechol (Sauret-Ignazi et al., 1996). In addition, high concentrations of 3-methylcatechol caused inhibition by substrate. Furthermore, this C12O is inhibited in the presence of phenol, diverse chlorophenols and fluorophenols (Sauret-Ignazi et al., 1996). The *phy-catA1* cluster may be controlled by PoxR, a XylR/NtrClike transcriptional regulator that acts as an activator in presence of phenol (Table S3), which also might be associated with proteins from the central metabolism, such as DmpF, MhpF and AtoA, as predicted by bioinformatics. It has been postulated that the presence of multiple BMMs in the same organism may lead to the formation of complex modularity generating new hybrids with new substrates specificity providing optimized metabolic pathways (Notomista et al., 2003; Cafaro et al., 2004).

Based on the experimental evidence presented in this report, we suggest that benzene is transformed into phenol via various routes catalyzed by three BMMs. Previous reports have demonstrated that the subsequent conversion of phenol into catechol is the limiting step in benzene aerobic degradation (Zhu et al., 2008). This would explain the phenol accumulation as an intermediate during the growth of CH34 on benzene (Figure 2B). The catechol formation was inferred by *meta*-cleavage of the catechol ring and subsequent formation of the colored compound 2-HMS. This suggests that one or more BMMs could be activated simultaneously during the process, thereby indicating the use of mixed degradation pathways, to generate a catabolic strategy that also seems to be deployed by *C. pinatubonensis* JMP134, *R. pickettii* PKO1, and *P. spadix* BD-a59 (Notomista et al., 2003; Tao et al., 2004; Perez-Pantoja et al., 2012; Choi et al., 2013). The *meta*-cleavage of the dihydroxylated ring indicates activity of a catechol-2,3-dioxygenase (C23O; TomB) that opens the catechol ring in *meta* position.

Results obtained by RT-qPCR showed a simultaneous expression of the three BMM-encoding gene clusters, and an increased expression of the gene encoding a C23O (*tomB*), HMSD (*tomC*), and HMSH (*tomD*) (Figure 3). The results obtained in gene expression analyses, at early and late exponential phases, are in accordance with our predictions based on gene sequence and organization (Figure 1, Table S2). A partial repression of the gene encoding the C12O, located downstream of the phenol-2-hydroxylase subunit genes (*phyZABCDE*), is in agreement with the results obtained via bioinformatic predictions for sigma 38 (*rpoS*) dependent gene expression. This sigma factor is predominately present in late exponential and stationary phases of growth (Tanaka et al., 1993; Jishage and Ishihama, 1995), which have also been observed in this study (Figure 3). As phenol inhibits C12O (Parales et al., 2008) it is expected that *catA* gene expression would be postponed until a later phase of growth, when phenol concentration, produced during the early growth phase, eventually decreases (Figure 2). Probably C23O activity from TomB might be temporarily preferred for phenol transformations in the early stages of growth. Furthermore, the capability to use either simultaneously or sequentially a C12O and a C23O may explain why some bacteria display a high versatility in their aromatic compounds degradation capability (Parales et al., 2008). For these reasons, in this report is postulated that strain CH34 possesses a mixed peripheral benzene degradation pathway deploying three functional BMMs. First, benzene is transformed into phenol catalyzed by one or both BMMs (T2MO and T4MO), followed by phenol-level mediated activation of the genes *phyZABCDE* encoding the subunits of a third BMM transforming the toxic phenol into catechol. This postulation is the core of the present work and is supported by the results presented here.

On the other hand, the results provide evidence that suggest a *meta*-cleavage of the catechol ring during the central pathway, catalyzed by a C23O (TomB). Conversely, an *ortho*-cleavage of the dihydroxylated ring, catalyzed by a C12O, will occurs at late stages of growth (Figure 3). These results are concordant with previous reports describing *C. metallidurans* CH34 as a versatile organism that rely on a complex transcriptional regulatory network for it to survive on highly diverse contaminated environments, *i.e*. soils or water with low nutrients levels and polluted with mixtures of metal ions (Monsieur et al., 2011).

One of the most important factors that affect microbial degradation of benzene in most microorganisms, it is the high toxicity even at low concentrations (Lovley, 1995; Sandrin and Maier, 2003). A good example is the model bacterial strain for benzene degradation, *P. putida* F1, which is only capable of growing at concentrations of up to 0.55 mM (Reardon et al., 2000). In this study, CH34 cells were able to grow in 5 mM benzene until turbidity of 0.6 at 600 nm (Figure 2A). Few bacterial strains are capable of growing on high concentrations of benzene. *Rhodococcus* sp. 33, tolerates up to benzene 1 mM as carbon and energy source (Paje et al., 1997). Such unusual capabilities of strain CH34 to use diverse aromatic compounds for growth (Table 1) and tolerate high benzene concentrations, suggests that a mixed degradation pathway might be beneficial for an organism’s robustness and versatility when it is faced to toxic levels of aromatic compounds. Additionally, strain CH34 possesses all the genes needed to catalyze oxidation of toluene and *o- m- p-* xylene isomers (Janssen et al., 2010; Mergeay and van Houdt, 2014), likely using the same benzene degradation pathway as proposed in Figure 4. This broad-range capacity also occurs in strains *C. pinatubonensis* JMP134, *R. pickettii* PKO1, and *P. spadix* BD-a59 (Shields et al., 1989; Parales et al., 2008; Perez-Pantoja et al., 2012; Choi et al., 2013). Nonetheless, further experimental studies are needed with strain CH34 to further characterize this metabolic pathway.

This study also revealed that *C. metallidurans* CH34 possesses the capability to grow on various aromatic compounds such as benzene, toluene, *o*-xylene, *p*-cymene, 3-hydroxybenzoate, 4-hydroxybenzoate, 3-hydroxyphenylacetate, 4-hydroxyphenylacetate, homogentisate, catechol, naphthalene, and 2-aminophenol as only carbon and energy source (Table 1). Previous studies have also reported the growth of *C. metallidurans* CH34 on the aromatic compounds benzoate, 4-hydroxybenzoate, phenol and tryptophan. In addition, strain CH34 has the capability to degrade phenylacetate and homogentisate (Mergeay and van Houdt, 2014). Our findings confirm that strain CH34 is capable of growing on either 3- or 4-hydroxybenzoate as a sole carbon and energy source (Table 1) and the genes that encode the complete benzoate degradation pathway, have been identified and found on the chromid of CH34 (Janssen et al., 2010; Perez-Pantoja et al., 2012; Mergeay and van Houdt, 2014). By bioinformatic studies, it is proposed that the enzymes responsible of the benzoate catabolic pathway are a benzoate-1,2-dioxygenase (*benABC* formerly Rmet_4882-Rmet_4884), a dihydroxybenzoate dehydrogenase (*benD* formerly Rmet_4885), and a catechol-1,2-dioxygenase, C12O (*catA2* formerly Rmet_4881), which *ortho*-cleaves the catechol ring and a subsequent conversion into intermediates of the tricarboxylic cycle (TCA) is catalyzed by three enzymes encoded by the *catBCD* locus (Table S4).

Degradation of aromatic compounds is inhibited in presence of heavy metals such as Pb(II) or Hg(II) (Said and Lewis, 1991; Benka-Coker and Ekundayo, 1998). In this study was demonstrated that *C. metallidurans* CH34 is able to degrade aromatic compounds, even in presence of toxic heavy metals such as Pb(II) or Hg(II) (Table 2). This strain is capable of growing on benzene (5 mM) as only carbon and energy source, in presence of Pb(II) (0.4 mM). In contrast to other strains, *C. metallidurans* CH34 is resistant to Hg(II) concentrations up to 0.08 mM (Table 2). Therefore, strain CH34 stands out among other benzene degradative strains to resist 0.08 highest concentrations of benzene (5 mM) in presence of toxic heavy metals. The derivative strain of CH34, *C. metallidurans* MSR33, which carries a natural plasmid IncP-1β (pTP6) providing additional set of *mer* genes and conferring an increased resistance to inorganic and organic mercury compounds (Smalla et al., 2006; Rojas et al., 2011), has the capability to reduce inorganic and organic forms of Hg(II) to metallic mercury, conferring to the strain MSR33 possible significantly improvements in terms of aromatic compounds degradation in presence of heavy metals (data not shown). Therefore, other *C. metallidurans* strains may be attractive catalysts for novel bioremediation applications in complex polluted environments *i.e*. where organisms have to cope with both heavy metals and aromatic compounds, such as mining sites.

This study has shown the aromatic compounds catabolic potential and versatility of the heavy metal resistant bacteria *C. metallidurans* CH34. Additionally, this report revealed the functionality of the benzene catabolic pathway that is active even in presence of mercury or lead. Strain CH34 is able to use diverse aromatic compounds as sole carbon and energy source, indicating active catabolic pathways for the degradation of benzene, toluene, *o-*xylene, *p-*cymene, 3-hydroxybenzoate, 4-hydroxybenzoate, 3-hydroxyphenylacetate, 4-hydroxyphenylacetate, homogentisate, catechol, naphthalene, and 2-aminophenol.

## Authors Contributions

Conceived and designed the experiments: FAM FC LAR MS.

Performed the experiments: FAM FC

Data analysis: FAM VM MS LAR

Contributed reagents/materials/analysis tools: MS LAR

Wrote the paper: FAM VM MS LAR

## Acknowledgments

The authors acknowledge Francisco Montero, Sebastian Fuentes and Paul Janssen for their helpful discussion and support with experimental analysis.

